# Graph convolutional networks for drug response prediction

**DOI:** 10.1101/2020.04.07.030908

**Authors:** Tuan Nguyen, Giang T.T. Nguyen, Thin Nguyen, Duc-Hau Le

## Abstract

**Background:** Drug response prediction is an important problem in computational personalized medicine. Many machine-learning-based methods, especially deep learning-based ones, have been proposed for this task. However, these methods often represent the drugs as strings, which are not a natural way to depict molecules. Also, interpretation (e.g., what are the mutation or copy number aberration contributing to the drug response) has not been considered thoroughly.

**Methods:** In this study, we propose a novel method, GraphDRP, based on graph convolutional network for the problem. In GraphDRP, drugs were represented in molecular graphs directly capturing the bonds among atoms, meanwhile cell lines were depicted as binary vectors of genomic aberrations. Representative features of drugs and cell lines were learned by convolution layers, then combined to represent for each drug-cell line pair. Finally, the response value of each drug-cell line pair was predicted by a fully-connected neural network. Four variants of graph convolutional networks were used for learning the features of drugs.

**Results:** We found that GraphDRP outperforms tCNNS in all performance measures for all experiments. Also, through saliency maps of the resulting GraphDRP models, we discovered the contribution of the genomic aberrations to the responses.

**Conclusion:** Representing drugs as graphs can improve the performance of drug response prediction.

**Availability of data and materials:** Data and source code can be downloaded at https://github.com/hauldhut/GraphDRP.

## 1 Introduction

**U**SING the right drug in the right dose at the right time is a goal in personalized medicine. Thus, estimating how each patient responds to a drug based on their biological characteristics (e.g., omics data) is important in biomedical research. However, patients’ drug response data is very limited and not well-structured. Indeed, there have been only a few studies on drug response for cancer patients gathered in TCGA [1]. This has formed a barrier to large-scale research on this topic.

Fortunately, large-scale projects on drug response for “artificial patients” (i.e., cell line), such as GDSC [2], CCLE [3] and NCI60 [4] have facilitated the development of computational methods for drug response prediction [5], [6], [7], [8]. Indeed, a DREAM challenge for drug sensitivity prediction was launched and with methods proposed by many research groups [9]. Most of them are machine learning-based, where different strategies for data and model integration were introduced. For example, multiple-kernel and multiple-task learning techniques were proposed to integrate various types of –omics data of cell lines and response data [10], [11]. Besides, ensemble learning strategies were used to integrate individual models [12], [13], [14]. In parallel, network-based methods relying on similarity networks (e.g., structural similarity between drugs and biological similarity between cell lines) and known drug-cell line responses [15], [16], [17] have been proposed. In addition, protein interaction and gene regulatory networks have also been used also used to predict drug response [18].

The machine learning-based methods have shown their ability in the data and model integration, thus drug response prediction was generally systematically approached. However, drugs and cell lines are often represented by predefined features such as structural features of drugs and -omics profiles of cell lines. As the number of cell lines is much smaller than the number of genes in -omics profiles of cell lines, thus some of the traditional machine learning-based methods often face the “small *n*, large *p*” problem. Consequently, this limits the prediction performance of traditional machine learning-based methods.

Deep learning is a state-of-the-art branch of machine learning for extracting a feature from complex data and making accurate predictions [19]. Recently, deep learning has been applied to drug discovery [20], [21]. It has achieved superior performance compared to traditional machine learning techniques in many problems in drug development such as visual screening [22], [23], drug-target profiling [24], [25], [26], [27], drug repositioning [28], [29]. Especially in drug response, deep learning is utilized to automatically learn genomic features of cell lines and the structural features of drugs to predict anticancer drug responsiveness [30], [31], [32], [33]. For example, the deep neural network is used in DeepDR [31] to predict the half-maximal inhibitory concentrations (IC50) or the convolutional neural network is utilized in tCNNS [33] and CDRScan [30] to extract the features of cell lines and drugs. In addition, in DeepDSC [32], a pre-trained stacked deep autoencoder is used to extract genomic features of cell lines from gene expression data and then combine with chemical features of compounds to predict response data. However, in these deep learning models, drugs are represented as strings which are not a natural presentation, thus the structural information of drugs may be lost.

Graph convolutional networks can learn representations of compound structures represented as molecular graphs [34]. For example, in GraphDTA [35], the drugs are presented as graphs where the edges are the bonding of atoms and the model achieves the best performance compared to other deep learning-based methods which represent drugs as strings in the task of drug-target binding affinity prediction. However, the graph neural network has not been employed yet [34] for drug response prediction. So it is promising to apply graph neural network to drug response prediction. In addition, although deep learning-based methods often achieve better prediction performance when compared to traditional machine learning-based methods, it is considered as a black-box approach because of not being interpretable. The saliency map [36] was introduced to visualize image features in classification task at first, now it plays an important role in various practical applications right from video surveillance [37] to traffic light detection [38]. In this research, this strategy can help evaluate the degree of genomics features such as aberration attributes to drug response prediction.

tCNNS was recently published and shown to be the state-of-the-art method among other deep learning-based methods [33]. We also tested traditional machine learning methods such as decision tree, gradient boosting but even tCNNS surpassed these methods. So in our work, we only compared our result directly with tCNNS.

In this study, we propose GraphDRP (Graph convolutional network for drug response prediction), a new neural network architecture capable of modeling drugs as molecular graphs to predict drug response on cell-line. We compared our method with the state-of-the-art, tCNNS [33], where drug molecules were represented as SMILES [39] strings. Experimental results indicate that our method achieves better performance in terms of root mean square error (RMSE) and Pearson correlation coefficient for all experiments. Also, by visualizing the resulting networks through saliency maps, we can discover the most significant genomic aberrations for the prediction of the response value. This suggests a novel way to interpret the result of deep learning models for drug response prediction.

## 2 Graph convolutional network for drug response prediction (GraphDRP)

The proposed model of drug response prediction is shown in Fig 1. The input data includes chemical information of drugs and genomic features of cell lines including mutations and copy number alternations (i.e., genomic aberration). For the drug features, the drugs represented in SMILES format [39] were downloaded from PubChem [40]. Then, RDKit, an open-source chemical informatics software [41], was used to construct a molecular graph reflecting interactions between the atoms inside the drug. Atom feature design from DeepChem [42] was used to describe a node in the graph. Each node contains five types of atom features: atom symbol, atom degree calculated by the number of bonded neighbors and Hydrogen, the total number of Hydrogen, implicit value of the atom, and whether the atom is aromatic. These atom features constituted a multi-dimensional binary feature vector [35]. If there exists a bond among a pair of atoms, an edge is set. As a result, an indirect, binary graph with attributed nodes was built for each input SMILES string. Several graph convolutional network models, including GCN [43], GAT [44], GIN [45] and combined GAT-GCN architecture [35], were used to learn the features of drugs. Following the graph neural network, a fully connected layer (FC layer) was also used to convert the result to 128 dimensions.

**Fig 1.**
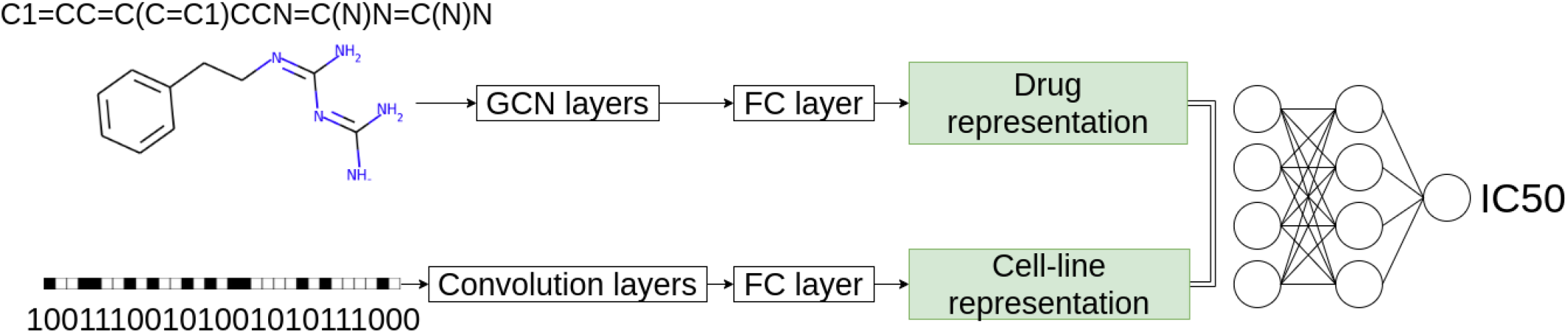
An illustration of GraphDRP. Each cell line was converted to one-hot format with a vector of 735 dimensions. Then 1D convolutional layers were applied three times to these features. After that, the fully connected (FC) layer was used to convert ConvNet results in 128 dimensions. Drug in SMILE string was converted to graph format. Then graph convolutional networks were used to learn the drug’s feature. Following the graph neural network, the fully connected layer was also used to convert the result to 128 dimensions. Finally, the two representations were then concatenated and put through two FC layers to predict the response.

In deep learning models, 1D convolutions are normally used for genomic features. We used the same approach as other models because 1D convolution with a large kernel has the ability to combine genomic abbreviations in the genomic features to make good predictions. In addition, 1D pooling was also used to reduce the size of input feature then 1D convolutions can learn abstract features from genomic features. The genomic features of cell lines were represented in one-hot encoding. 1D convolutional neural network (CNN) layers were used to learn latent features on those data. Then the output was flattened to 128 dimension vector of cell line representation.

Finally, the 256-dimension vector, the combination of drug’s feature and cell line’s feature was put through two fully-connected layers with the number of nodes 1024 and 256 respectively, before predicting the response.

CNNs have recently achieved success in computer vision [46], [47] and natural language processing [48], [49], which motivates the use of convolutional neural networks to graph structures. Similar to the use of CNN with image, CNN in the graph also has two main layers: convolutional layer and pooling layer. While convolutional layer is for learning receptive fields in graphs whose data points are not arranged as Euclidean grids, pooling layer is for down-sampling a graph [35]. Graph convolutional network (GCN) is well-fitted for the drug response because the drug molecular itself is represented in the form of a graph. In order to evaluate the effectiveness of graph-based models, we investigated several graph convolutional models, including GCN [43], GAT [44], GIN [45] and combined GAT-GCN architecture [35]. The details of each GCN architecture are described as follows.

### 2.1 Graph Convolutional Networks (GCN)

Formally, a graph for a given drug *G* = (*V, E*) was stored in the form of two matrices, including feature matrix *X* and adjacency matrix *A. X* ∈ *R*^*N×F*^ consists of N nodes in the graph and each node is represented by *F* -dimensional vector. *A* ∈ *R*^*N×N*^ displays the edge connection between nodes. The original graph convolutional layer takes two matrices as input and aims to produce a node-level output with *C* features each node. The layer is defined as:

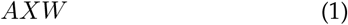

where *W* ∈ *R*^*F ×C*^ is the trainable parameter matrix. However, there are two main drawbacks. First, for every node, all feature vectors of all neighboring nodes were summed up but not the node itself. Second, matrix A was not normalized, so the multiplication with A will change the scale of the feature vector. GCN model [43] was introduced to solve these limitations by adding identity matrix to A and normalizing A. Also, it was found that symmetric normalization achieved more interesting results. The GCN layer is defined by [43] as

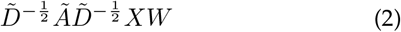

where *Ã* is the graph adjacency matrix with added self loop, 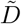 is the graph diagonal degree matrix.

In our GCN-based model, three consecutive GCN layers were utilized and ReLU function was applied after each layer. A global max pooling layer was added right after the last GCN layer to learn the representation vector of whole graph and then combine with the representation of cell-line to make the prediction of response value.

### 2.2 Graph Attention Networks (GAT)

Self-attention technique has been shown to be self-sufficient for state-of-the-art-level results on machine translation [50]. Inspired by this idea, we used self-attention technique in graph convolutional network in GAT [44]. We adopted a graph attention network (GAT) in our model. The proposed GAT architecture was built by stacking a *graph attention layer*. The GAT layer took the node feature vector **x**, as input, then applied a linear transformation to every node by a weight matrix **W**. Then the *attention coefficients* is computed at every pair of nodes that the edge exists. The coefficients between node *i* and *j* were computed as

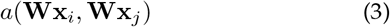

This value indicates the importance of node *j* to node *i*. These *attention coefficients* were then normalized by applying a soft-max function. Finally, the output features for each node was computed as

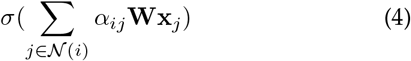

where *σ*(.) is a non-linear activation function and *α*_*ij*_ are the normalized *attention coefficients*.

In our model, we used two GAT layers, activated by a ReLU function, then a global max pooling layer was followed to obtain the graph representation vector. In details, for the first GAT layer, we used *multi-head-attentions* with 10 heads, and the number of output features was equal to the number of input features. The number of output features of the second GAT was set to 128, similar to cell-line representation vector.

### 2.3 Graph Isomorphism Network (GIN)

We adopted a recently proposed graph learning method, namely Graph Isomorphism Network (GIN) [45] in our model. It is theoretically proven to achieve maximum discriminative power among GNNs [45]. Specifically, the node feature was updated by multi layer perceptron (MLP) as

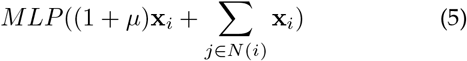

while *µ* is either a learnable parameter or a fixed scalar, **x** is the node feature vector, and *N* (*i*) is the set of nodes neighbor to *i*.

In our model, five GIN layers with 32 features were stacked to build GIN architecture. For each layer, batch normalization layer was used, then ReLU activation function was applied to learn the nonlinear mapping function. Similar to GAT architectures, a global max pooling layer was added to aggregate a graph representation vector.

### 2.4 Combined graph neural network (GAT&GCN)

A combination of GAT [44] and GCN [43] was also proposed to learn graph features [35]. At first the GAT layers learned to combine nodes in attention manner, so after GAT layers these features in each node were abstract and contained high-level information of the graph. Finally, GCN layers were used to learn convolved features to combine these abstract nodes to make final prediction.

## 3 Model Interpretation: GenomicAberration Contribution Using Saliency Map

Given a drug-cell line pair, saliency value was defined by using the idea of the saliency map [36] to measure the importance of each genomic aberration to the prediction of response value (Y). In our proposed model, each drug (D) was represented by a graph, meanwhile, each cell-line (C) was displayed by a binary vector of 735 dimensions, with each value indicates whether or not the cell-line had a specific genomic aberration.

f is the whole deep learning model function.

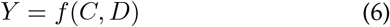

Then saliency value (S) was defined as the gradient of cell-line with respect to predicted response as follows:

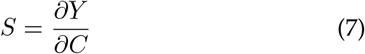

This saliency value has the same size as cell-line vector. The higher value indicates the more important of genomic aberration that was encoded in this position.

## 4 Experimental Setting

### 4.1 Datasets

Large-scale drug sensitivity screening projects such as CCLE [3] and GDSC [2] generated not only –omics but also drug response data for anti-cancer drugs on thousands of cell lines. The –omics data includes gene expression (i.e., transcriptomic data), which indicates an amount of RNAs transcribed from DNA and thus amount of translated proteins in a cell. Therefore, the expression level of a gene indicates the activity level of a gene in a certain state (e.g., diseased or normal) in a cell. In addition, the –omics data also implies genomic aberrations such as mutations and copy number variations (CNVs) in genome. Meanwhile, drug response is a measure of drug efficiency to inhibit the vitality of cancer cells. More specifically, cell lines are cultured and treated with different doses of drugs. Finally, either an IC50 value, which indicates the dose of a particular drug needed to inhibit the biological activity by half, or an AUC (area under dose-response curve) value is used as a response measure of a particular drug.

GDSC is the largest database of drug sensitivity for cell lines. Indeed, there are 250 drugs tested on 1,074 cell lines in that database, meanwhile only 24 drugs were tested on 504 cell lines in CCLE. Thus, we selected GDSC version 6.0 as the benchmark dataset for this study. Following the same procedure as in tCCN [33], after preprocessing, 223 drugs and 948 cell lines were finally selected. A total of 172,114 (81.4%) drug-cell line pairs were tested with response values, and the remaining (18.6%) of pairs were missing. Similarly, the response values in terms of IC50 were also normalized in a range (0,1) as in [51]. In addition, at the input stage, a cell line was described by a binary vector of 735 dimensions, where 1 or 0 indicate whether a cell line has or has not a genomic aberration respectively. Meanwhile, drugs were represented in canonical SMILES format [39].

### 4.2 Experimental design

In this section, the performance of our model is demonstrated through three experiments: performance comparison, prediction of unknown drug-cell line response and investigation of genomic aberration contribution to the response. To compare to previous studies, the same setting was used for performance comparison. Several graph models including GCN, GIN, GAT, GCN GAT were used to learn the representation of the drug. The prediction of unknown response pairs and the contribution of genomic aberrations by saliency map help to interpret the proposed model. Initially, we chose the hyperparameter values based on previous work. Then we tuned a lot of parameters such as learning rate, batch size to achieve the better result. Detail experiments are described below.

#### 4.2.1 Performance comparison

##### Mixed test

This experiment evaluates the performance of models in known drug-cell line pairs. Of all 211,404 possible drug-cell line pairs, GDSC provides the response for 172,114 pairs [33]. The data was shuffled before splitting to help the model remain general and reduce overfitting. The known pairs are split into 80% as the training set, 10% as the validation set and 10% as the testing set. While the validation set was used to modify the hyperparameter of the model in the training phase, the testing set was used to evaluate the performance of the model.

##### Blind test

In the previous experiment, a drug which had appeared in the testing set might also appear in the training phase. However, we sometimes need to predict the response of a new drug, for example, a newly invented one. This experiment was designed to evaluate the prediction performance of unseen drugs. Drugs were constrained from existing in training and testing at the same time. Of 90% (201/223) drugs, their IC50 values were randomly selected for training, including 80% drugs for the training set and 10% drugs for the validation set. The remaining set, 10% (22/223) drugs were used as the testing set.

Similarly, it is sometimes required to make predictions for a new cell-line that are not in the training phase. So we also did an experiment to test the prediction of unseen cell-lines. Cell-lines were constrained from existing in training and testing at the same time. A total of 90% (891/990) cell-lines were randomly selected and their IC50 values were kept for the training phase. The remaining, 10% (99/990) cell-lines, was used as the testing set.

#### 4.2.2 Prediction of unknown drug-cell line response

This experiment aims at predicting missing drug-cell line response. The best pre-trained model in the mixed test experiment was used to predict missing pairs in GDSC dataset. Then we selected top 10 drugs that had the lowest and highest IC50 values to further investigate whether drugs having lower IC50 are more effective to the treatment of cancer and whether the ones having higher IC50 are less effective.

#### 4.2.3 Investigation of genomic aberration contribution to the response

Given a pair of drug and cell-line, the saliency value is calculated by taking derivative of predicted response with respect to the cell-line, in one-hot vector format. To do this experiment, we chose a drug with the lowest average IC50 among all drugs in the prediction of unknown drug-cell line response. Then we selected three cell-lines that have the lowest IC50 values to that drug. Then, for each pair, we calculated the saliency value then took ten most important genomic aberrations. Eventually, we found evidence to support the contribution of the genomic aberrations to the response of that drug on the selected cell-lines.

### 4.3 Performance Evaluation

Two metrics were adopted to measure the performance of models: root mean square error (RMSE) and Pearson correlation coefficient (*CC*_*p*_).

Given *O* the ground-truth, *Y* the predicted value, n is the number of samples, *o*_*i*_ is a ground-truth of *i*^*th*^ sample, *y*_*i*_ is the predicted value of *i*^*th*^ sample. RMSE measures the difference of them

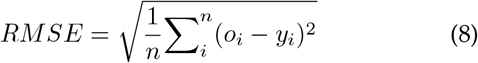

where *n* is the number of data points.

Pearson correlation coefficient measures how strong a relationship is between two variables. Let the standard deviation of O and Y be *σ*_*O*_, *σ*_*Y*_ respectively, *CC*_*p*_ is defined as

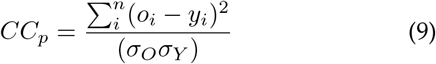

The lower RMSE, the better the model is. Whereas, the higher *CC*_*p*_, the better the model is.

## 5 Results and Discussion

### 5.1 Performance comparison

Tables 1, 2 & 3 present the prediction performance in terms of *CC*_*p*_ and RMSE for different experiments by the baseline (tCNNS [33]) and our proposed method.

**TABLE 1.** Performance comparison in terms of *CC*_*p*_ coefficient and RMSE on the GDSC dataset in the mixed test experiment. The best performance is in bold.

| Method | | $CC_p$ | RMSE |
| --- | --- | --- | --- |
| tCNNS [33] |  | 0.9160 | 0.0284 |
| GraphDRP | GCN | 0.9216 | 0.0259 |
|  | GIN | **0.9310** | 0.0244 |
|  | GAT | 0.9270 | 0.0250 |
|  | GCN_GAT | 0.9308 | **0.0243** |

**TABLE 2.** Performance comparison in terms of *CC*_*p*_ and RMSE on the GDSC dataset in the blind test with the unseen drug experiment. The best performance is in bold.

| Method | | $CC_p$ | RMSE |
| --- | --- | --- | --- |
| tCNNS [33] |  | 0.0617 | 0.0680 |
| GraphDRP | GCN | **0.3241** | **0.0542** |
|  | GIN | 0.0481 | 0.0602 |
|  | GAT | 0.2751 | 0.0616 |
|  | GCN_GAT | 0.1683 | 0.0610 |

**TABLE 3.** Performance comparison in terms of *CC*_*p*_ and RMSE on the GDSC dataset in the blind test with unseen cell-line experiment. The best performance is in bold.

| Method | | $CC_p$ | RMSE |
| --- | --- | --- | --- |
| tCNNS [33] |  | 0.3490 | 0.0576 |
| GraphDRP | GCN | 0.8399 | 0.0363 |
|  | GIN | **0.8460** | **0.0358** |
|  | GAT | 0.8312 | 0.0380 |
|  | GCN_GAT | 0.8402 | 0.0362 |

#### Mixed test

In this experiment, we evaluate and compare the prediction performance of GraphDRP with tCNNS. RMSE and *CC*_*p*_ were calculated for both methods based on the same benchmark dataset and settings. The performance of the two methods is shown in Table 1. It is obvious that our model GraphDRP outperforms tCNNS for all graph convolutional networks. tCNNS achieved a RMSE of 0.0284 and a *CC*_*p*_ of 0.9160, meanwhile the worse RMSE and *CC*_*p*_ in our models were 0.0259 and 0.9216, respectively. GraphDRP achieved the best RMSE (0.0243) with GIN model and the best *CC*_*p*_ (0.9308) with GCN GAT model. For RMSE, GIN obtained the second best result (0.0244), which was just a little smaller than the best result (0.0243). Thus, we considered it the best model in this experiment.

#### Blind test

In the mixed test experiment, one drug should have been presented in both training and testing sets. However, it was more challenging to predict the response of unseen drugs/cell-lines. So in this experiment, drugs/cell-lines in the testing stage were not present in the training stage.

Table 2 shows the prediction performance for the blind test with unseen drugs. We observed that our proposed models, for all kinds of convolution graphs, achieved better RMSE than tCNNS. In particular, the GCN gained the best RMSE of 0.0542. Meanwhile, for *CC*_*p*_, except for GIN, other three graph-based methods gained better performance when compared to tCNNS. Particularly, GCN-based gained a five-fold increase, 0.3241 versus 0.0617 compared to tCNNS in terms of *CC*_*p*_ and it was the best method in terms of both *CC*_*p*_ and RMSE in this experiment.

Similar to the drug blind test, this experiment evaluates the performance of the model on unseen cell-lines. Cell-lines are prevented from existing in the training and testing phase at the same time. In the prediction of response for unknown cell-line, our proposed methods achieved better performance in terms of both RMSE and *CC*_*p*_ than tCNNS for all kinds of GCN. Particularly, GIN method gained the best *CC*_*p*_ at 0.8460 and the best RMSE at 0.0358.

For both drug and cell-line blind test, the prediction performances were not good as that in mixed test experiment for all models. This indicates that it is harder to predict the drug-cell line response for unseen drugs or unseen cell-lines. Interestingly, we observed that the performance of predicting response for unseen cell-line (i.e., average values were 0.8393 (±0.0061) for *CC*_*p*_ and 0.0366 (±0.001) for RMSE) is better than that for unseen drug (i.e., average values are 0.2039 (±0.1226) and 0.059 (±0.0034) for *CC*_*p*_ and RMSE respectively).

We tested several graph models including GCN, GIN, GAT, GCN GAT for learning the representation of drugs. It is clearly shown that our models outperformed the tCNNS in all experiments. This is because, in tCNNS, the SMILE string format used for drug was not natural representation. While in our model, the graph convolutional networks were used to extract information from graph representation of drugs, so the performance was better.

Amongst three experiments, GIN achieved the best prediction performance in terms of both RMSE and *CC*_*p*_ in the mixed test and the blind test with unseen cell-lines. It unleashed the potential of GIN in graph representation, partly supporting the claim in [45] that GIN is among the most powerful GCNs. However, it is noticeable that in these two experiments, all drugs were observed in both training set and validation set so that the GIN model could learn the features of drugs. However, in the blind test for unseen drugs, the drugs in the validation sets did not appear in the training set. As a result, the GIN model did not perform well in this experiment.

### 5.2 Prediction of unknown drug-cell line response

In this experiment, the best model trained on the mixed test experiment (i.e., GIN) was used to predict the response for 39,290 missing pairs. Figure 2 shows top 10 drugs that have the highest and lowest predicted IC50. Interestingly, top 3 drugs that have the highest and lowest IC50 values are the same result as in tCNNS. It is shown that *Bortezomib* achieveed the lowest IC50, which means it is the most sensitive drug for anti-cancer. Indeed, it was reported that *Bortezomib* had differential cellular and molecular effects in human breast cancer cells [52]. Also, it had a wide range of applications in antitumor activity [53]. The second most effective drug for anti-cancer in this experiment was *Epothilone B*, which acted by blocking cell division through its interaction with tubulin [54]. In contrast, *AICA Ribonucleotide* and *Phenformin* had the highest IC50, which means cancers were less sensitive to these drugs. Indeed, while AICA Ribonucleotide has been used clinically to treat and protect against cardiac ischemic injury [55], Phenformin is an antidiabetic drug from the biguanide class [33]. Thus, they were not designed to cure cancer.

**Fig 2.**
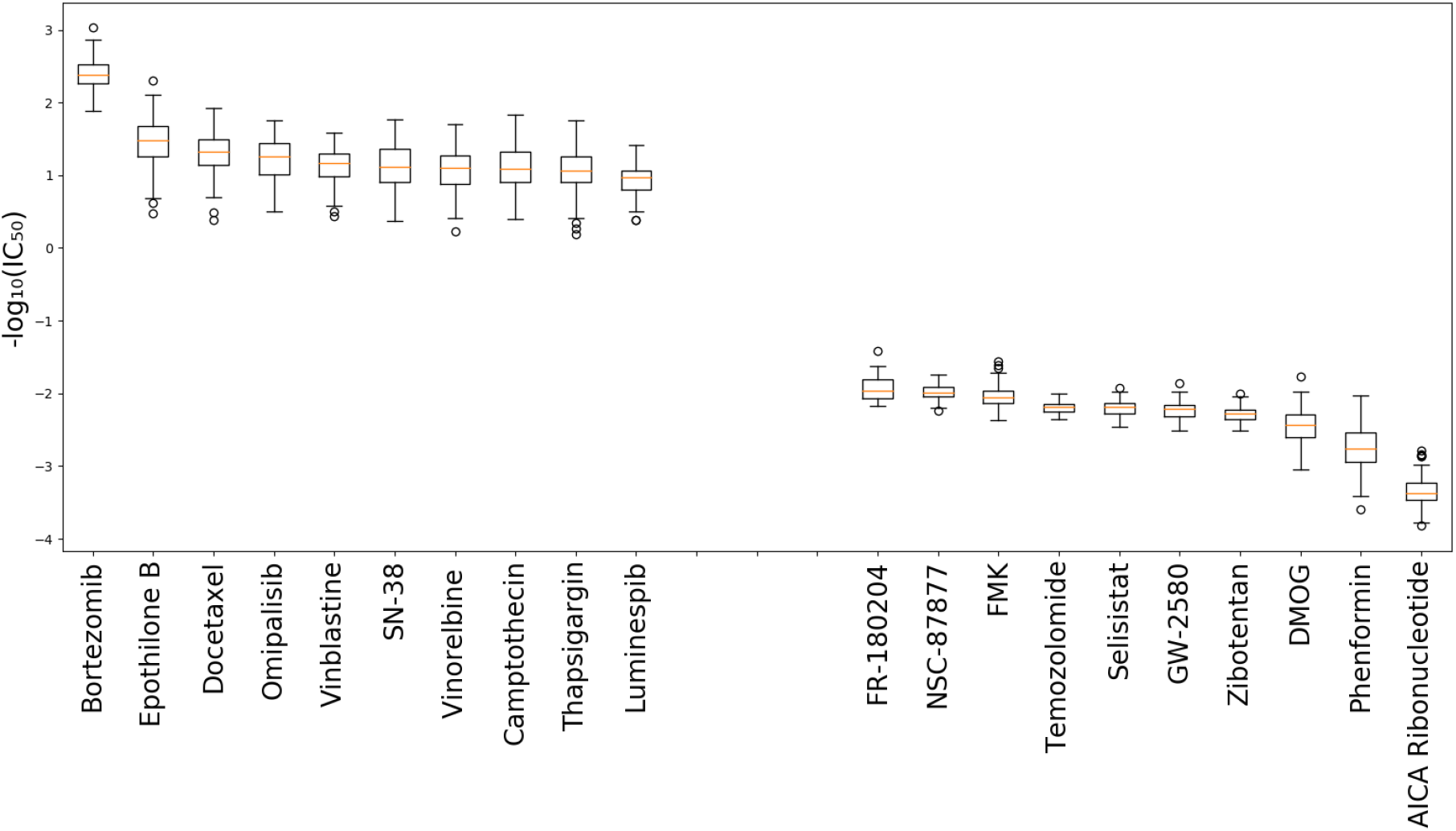
The boxplot of predicted missing value for top 10 drugs that have highest IC50 values and top 10 drugs that have lowest IC50 values.

Those evidence show that cancers are more sensitive to drugs having low IC50, meanwhile, cancers are less sensitive to those having high IC50. The results also indicate that our model is potential to the prediction of untested drug-cell line pairs.

### 5.3 Investigation of genomic aberration contribution to the response

*Bortezomib*, the most sensitive drug, was chosen in this experiment to further investigate the contribution of genomic aberrations to the response on cell-lines. Thus three cell-lines that had the lowest IC50 with that drug were taken to do the experiment. Saliency value of each genomic aberration is considered as the degree of their contribution to the response. Table 4 shows ten most contributed genomic aberrations for the three cell-lines. Some evidence was found from literature to support their contribution. Indeed, a study [56] showed that TP53 mutation (i.e., a mutation in cell line EW-3) was targeted by *Bortezomib*. In addition, the combination of *Bortezomib*, standard chemotherapy, and HDAC inhibition is currently being evaluated in clinical trials for MLL mutation (i.e, a mutation in cell line NCI-H748) [57]. It means that the model paids more attention to the genomic aberrations that are related to the corresponding drug. As a result, the GraphDRP model could be interpretable as it could identify which genomic aberrations in cell-lines were mainly responsible for the response of a particular drug.

**TABLE 4.** Ten most important genomic aberrations, sorted by the saliency, in the prediction of Bortezomib against NCI-H748, SW684 and EW-3 cell-lines

| NCI-H748 |  | SW684 |  | EW-3 |  |
| --- | --- | --- | --- | --- | --- |
| Aberration | Saliency | Aberration | Saliency | Aberration | Saliency |
| EWSR1-FLI1_mut | 0.0131 | cnaPANCAN84 | 0.0099 | cnaPANCAN257 | 0.0123 |
| NSD1_mut | 0.0059 | cnaPANCAN247 | 0.0028 | cnaPANCAN120 | 0.0058 |
| cnaPANCAN144 | 0.0037 | cnaPANCAN13 | 0.0026 | ASXL1_mut | 0.0047 |
| cnaPANCAN108 | 0.0036 | cnaPANCAN384 | 0.0023 | TP53_mut | 0.0046 |
| MAP3K4_mut | 0.0027 | cnaPANCAN383 | 0.0019 | ASH1L_mut | 0.004 |
| PCDH18_mut | 0.0027 | BMPR2_mut | 0.0019 | NR4A2_mut | 0.0031 |
| cnaPANCAN247 | 0.0027 | cnaPANCAN22 | 0.0018 | TP53BP1_mut | 0.003 |
| SRGAP3_mut | 0.0025 | cnaPANCAN211 | 0.0018 | cnaPANCAN238 | 0.0027 |
| MLL_mut | 0.0023 | cnaPANCAN143 | 0.0018 | cnaPANCAN247 | 0.0025 |
| cnaPANCAN146 | 0.0021 | cnaPANCAN377 | 0.0017 | cnaPANCAN103 | 0.0025 |

## 6 Conclusions and Discussion

In this study, we proposed a novel method for drug response prediction, called GraphDRP. In our model, drug molecules were presented as graphs instead of strings, cell-lines were encoded into one-hot vector format. Then graph convolutional layers were used to learn the features of compounds and 1D convolutional layers were used to learn cell-line representation. After that the combination of drug and cell-line representation was used to predict IC50 value. Four variants of graph neural networks including GCN, GAT, GIN and combination of GAT&GCN were used for learning features of drugs. We compared our method with state-of-the-art one, tCNNS [33], where drug molecules were represented as SMILES strings.

Experimental results indicate that our method achieves better performance in terms of both root mean square error and Pearson correlation coefficient. The performance suggests that representing drugs in graphs is more suitable than in strings since it conserves the nature of chemical structures of drugs. Furthermore, the responses of missing drug-cell line pairs in GDSC dataset were predicted and analyzed. We figured out that *Bortezomib* and *Epothilone B* have the lowest IC50 values and we found the evidence showing that some types of cancer are sensitive to these drugs. Similarly, we also found that cancers are less sensitive to drugs having the highest IC50 values. It means that the model actually learns from data and has a potential to predict the response of new drug-cell line pairs. Also, through saliency maps, we discovered ten most important genomic aberrations of the three cell-lines having lowest IC50s to that drug and seek their contribution to the sensitivity of that drug. This technique suggests a novel way to interpret the result of deep learning model in drug response.

Since cell lines in the same tissue types will share similar genetic information, in the future, we will split the cell lines based on different tissue types to demonstrate how the performance varies as cell line similarity decreases.

Because all drugs are screened at a certain concentration, if this concentration is low, that does not necessarily make it a good cancer drug. Rather, a drug that is particularly toxic to a cancer cell line compared to non-cancer cells is a good drug for this cell line. However, this non-cancer toxicity is not measured in the GDSC panel, and hence only looking at a low IC50 value is not sufficient. In this study, we only focused on improving the prediction of IC50 values by using GCNs. Also, in our study, we concentrated on improving drug response prediction by extracting drug features from their graph representation by GCN; thus, we used the same dataset with the tCNNS study, which learned drug’s features from their string format. In addition, only genomic data of cell lines was used in our study. Therefore, in future work, we will additionally use other -omics data such as gene expression, since it is proven to support drug response prediction [58] [59] [60].

---

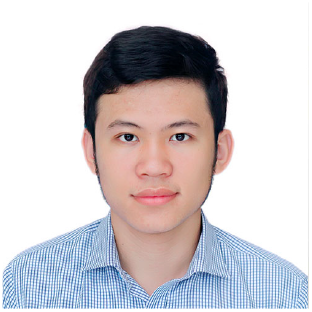

**Tuan Nguyen** is a data scientist at Big Data Institute, Vingroup. He received his MSc degree from University of Bristol in 2019 in the area of machine learning. His current research topic includes application of deep learning in medicine and the application of GAN.

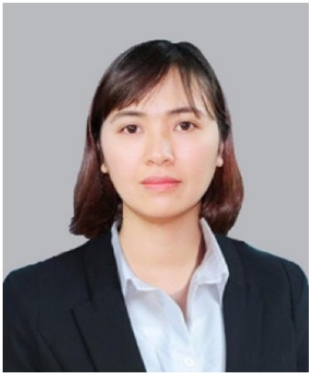

**Giang T**.**T. Nguyen** is a PhD Candidate in Information System at Posts and Telecommunications Institute of Technology. She received BSc and MSc in Computer Science at Hanoi University of Science and Technology. Her current research interests include Bioinformatics and Computational biology.

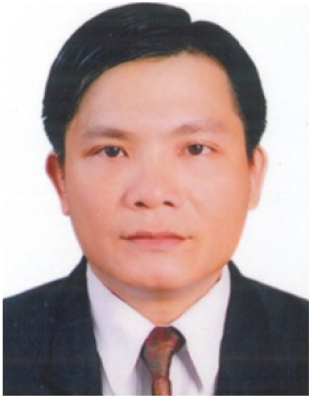

**Thin Nguyen** is a Senior Research Fellow with the Applied Artificial Intelligence Institute (A2I2), Deakin University, Australia. He graduated with a PhD in Computer Science from Curtin University, Australia in the area of machine learning and social media analytics. His current research topic is strongly inter-disciplinary, bridging large-scale data analytics, pattern recognition, genetics and medicine. His research direction is to utilize artificial intelligence to discover functional connections between drugs, genes and diseases.

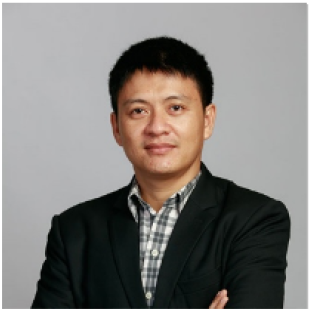

**Duc-Hau Le** received his Ph.D. degree in 2012 with research interests in bioinformatics and computational biology. So far, he has published many papers on prestigious journals. He is now leading the Department of Computational Biomedicine, Vingroup Big Data Institute in Vietnam, which aims to develop/apply computational methods to build and analyze high-throughput biomedical data in order to improve the screen, diagnosis and treatment of complex diseases. Specially, he is the principal investigator of the biggest genome project in Vietnam (i.e., building databases of genomic variants for Vietnamese population).

## References

[1] J. N. Weinstein, E. A. Collisson, G. B. Mills, K. R. M. Shaw, B. A. Ozenberger, K. Ellrott, I. Shmulevich, C. Sander, J. M. Stuart, C. G. A. R. Network et al., “The Cancer Genome Atlas Pan-Cancer analysis project,” Nature Genetics, vol. 45, no. 10, p. 1113, 2013.

[2] W. Yang, J. Soares, P. Greninger, E. J. Edelman, H. Lightfoot, S. Forbes, N. Bindal, D. Beare, J. A. Smith, I. R. Thompson et al., “Genomics of Drug Sensitivity in Cancer (GDSC): a resource for therapeutic biomarker discovery in cancer cells,” Nucleic Acids Research, vol. 41, no. D1, pp. D955–D961, 2012.

[3] J. Barretina, G. Caponigro, N. Stransky, K. Venkatesan, A. A. Margolin, S. Kim, C. J. Wilson, J. Lehár, G. V. Kryukov, D. Sonkin et al., “The Cancer Cell Line Encyclopedia enables predictive modelling of anticancer drug sensitivity,” Nature, vol. 483, no. 7391, pp. 603– 607, 2012.

[4] R. H. Shoemaker, “The NCI60 human tumour cell line anticancer drug screen,” Nature Reviews Cancer, vol. 6, no. 10, pp. 813–823, 2006.

[5] F. Azuaje, “Computational models for predicting drug responses in cancer research,” Briefings in Bioinformatics, vol. 18, no. 5, pp. 820–829, 2017.

[6] J. Chen and L. Zhang, “A survey and systematic assessment of computational methods for drug response prediction,” Briefings in Bioinformatics, 01 2020, bbz164. [Online]. Available: https://doi.org/10.1093/bib/bbz164

[7] M. J. Garnett, E. J. Edelman, S. J. Heidorn, C. D. Greenman, A. Dastur, K. W. Lau, P. Greninger, I. R. Thompson, X. Luo, J. Soares et al., “Systematic identification of genomic markers of drug sensitivity in cancer cells,” Nature, vol. 483, no. 7391, pp. 570–575, 2012.

[8] I. S. Jang, E. C. Neto, J. Guinney, S. H. Friend, and A. A. Margolin, “Systematic assessment of analytical methods for drug sensitivity prediction from cancer cell line data,” in Biocomputing. World Scientific, 2014, pp. 63–74.

[9] J. C. Costello, L. M. Heiser, E. Georgii, M. Gönen, M. P. Menden, N. J. Wang, M. Bansal, P. Hintsanen, S. A. Khan, J.-P. Mpindi et al., “A community effort to assess and improve drug sensitivity prediction algorithms,” Nature Biotechnology, vol. 32, no. 12, p. 1202, 2014.

[10] M. Gönen and A. A. Margolin, “Drug susceptibility prediction against a panel of drugs using kernelized Bayesian multitask learning,” Bioinformatics, vol. 30, no. 17, pp. i556–i563, 2014.

[11] D.-H. Le and D. Nguyen-Ngoc, “Multi-task regression learning for prediction of response against a panel of anti-cancer drugs in personalized medicine,” in Proceedings of the International Conference on Multimedia Analysis and Pattern Recognition (MAPR). IEEE, 2018, pp. 1–5.

[12] K. Matlock, C. De Niz, R. Rahman, S. Ghosh, and R. Pal, “Investigation of model stacking for drug sensitivity prediction,” BMC Bioinformatics, vol. 19, no. 3, p. 71, 2018.

[13] M. Tan, O.F. özgül, B. Bardak, I. Ekśioğlu, and S. Sabuncuoğlu, “Drug response prediction by ensemble learning and drug-induced gene expression signatures,” Genomics, vol. 111, no. 5, pp. 1078–1088, 2019.

[14] Q. Wan and R. Pal, “An ensemble based top performing approach for NCI-DREAM drug sensitivity prediction challenge,” PLoS ONE, vol. 9, no. 6, 2014.

[15] D.-H. Le and V.-H. Pham, “Drug response prediction by globally capturing drug and cell line information in a heterogeneous network,” Journal of Molecular Biology, vol. 430, no. 18, pp. 2993–3004, 2018.

[16] G. T. Nguyen and D.-H. Le, “A matrix completion method for drug response prediction in personalized medicine,” in Proceedings of the International Symposium on Information and Communication Technology, 2018, pp. 410–415.

[17] N. Zhang, H. Wang, Y. Fang, J. Wang, X. Zheng, and X. S. Liu, “Predicting anticancer drug responses using a dual-layer integrated cell line-drug network model,” PLoS Computational Biology, vol. 11, no. 9, 2015.

[18] T. Turki and Z. Wei, “A link prediction approach to cancer drug sensitivity prediction,” BMC Systems Biology, vol. 11, no. 5, p. 94, 2017.

[19] Y. LeCun, Y. Bengio, and G. Hinton, “Deep learning,” Nature, vol. 521, no. 7553, pp. 436–444, 2015.

[20] I. I. Baskin, D. Winkler, and I. V. Tetko, “A renaissance of neural networks in drug discovery,” Expert Opinion on Drug Discovery, vol. 11, no. 8, pp. 785–795, 2016.

[21] A. Lavecchia, “Deep learning in drug discovery: opportunities, challenges and future prospects,” Drug Discovery Today, 2019.

[22] A. Gonczarek, J. M. Tomczak, S. Zareba, J. Kaczmar, P. Dabrowski, and M. J. Walczak, “Interaction prediction in structure-based virtual screening using deep learning,” Computers in Biology and Medicine, vol. 100, pp. 253–258, 2018.

[23] J. C. Pereira, E. R. Caffarena, and C. N. dos Santos, “Boosting docking-based virtual screening with deep learning,” Journal of Chemical Information and Modeling, vol. 56, no. 12, pp. 2495–2506, 2016.

[24] M. Karimi, D. Wu, Z. Wang, and Y. Shen, “DeepAffinity: interpretable deep learning of compound–protein affinity through unified recurrent and convolutional neural networks,” Bioinformatics, vol. 35, no. 18, pp. 3329–3338, 2019.

[25] H. öztürk, A. özgür, and E. Ozkirimli, “DeepDTA: deep drug– target binding affinity prediction,” Bioinformatics, vol. 34, no. 17, pp. i821–i829, 2018.

[26] L. Wang, Z.-H. You, X. Chen, S.-X. Xia, F. Liu, X. Yan, Y. Zhou, and K.-J. Song, “A computational-based method for predicting drug– target interactions by using stacked autoencoder deep neural network,” Journal of Computational Biology, vol. 25, no. 3, pp. 361– 373, 2018.

[27] M. Wen, Z. Zhang, S. Niu, H. Sha, R. Yang, Y. Yun, and H. Lu, “Deep-learning-based drug–target interaction prediction,” Journal of Proteome Research, vol. 16, no. 4, pp. 1401–1409, 2017.

[28] A. Aliper, S. Plis, A. Artemov, A. Ulloa, P. Mamoshina, and A. Zhavoronkov, “Deep learning applications for predicting pharmacological properties of drugs and drug repurposing using transcriptomic data,” Molecular Pharmaceutics, vol. 13, no. 7, pp. 2524–2530, 2016.

[29] X. Zeng, S. Zhu, X. Liu, Y. Zhou, R. Nussinov, and F. Cheng, “deepDR: a network-based deep learning approach to in silico drug repositioning,” Bioinformatics, vol. 35, no. 24, pp. 5191–5198, 2019.

[30] Y. Chang, H. Park, H.-J. Yang, S. Lee, K.-Y. Lee, T. S. Kim, J. Jung, and J.-M. Shin, “Cancer drug response profile scan (CDRscan): a deep learning model that predicts drug effectiveness from cancer genomic signature,” Scientific Reports, vol. 8, no. 1, pp. 1–11, 2018.

[31] Y.-C. Chiu, H.-I. H. Chen, T. Zhang, S. Zhang, A. Gorthi, L.-J. Wang, Y. Huang, and Y. Chen, “Predicting drug response of tumors from integrated genomic profiles by deep neural networks,” BMC Medical Genomics, vol. 12, no. 1, p. 18, 2019.

[32] M. Li, Y. Wang, R. Zheng, X. Shi, F. Wu, J. Wang et al., “DeepDSC: A deep learning method to predict drug sensitivity of cancer cell lines,” IEEE/ACM Transactions on Computational Biology and Bioinformatics, 2019.

[33] P. Liu, H. Li, S. Li, and K.-S. Leung, “Improving prediction of phenotypic drug response on cancer cell lines using deep convolutional network,” BMC Bioinformatics, vol. 20, no. 1, p. 408, 2019.

[34] D. Baptista, P. G. Ferreira, and M. Rocha, “Deep learning for drug response prediction in cancer,” Briefings in Bioinformatics, 01 2020, bbz171. [Online]. Available: https://doi.org/10.1093/bib/bbz171

[35] T. Nguyen, H. Le, T. P. Quinn, T. Le, and S. Venkatesh, “Predicting drug–target binding affinity with graph neural networks,” bioRxiv, p. 684662, 2020.

[36] K. Simonyan, A. Vedaldi, and A. Zisserman, “Deep inside convolutional networks: Visualising image classification models and saliency maps,” arXiv preprint arXiv:1312.6034, 2013.

[37] A. Sobral, T. Bouwmans, and E.-h. ZahZah, “Double-constrained RPCA based on saliency maps for foreground detection in automated maritime surveillance,” in Proceedings of the IEEE International Conference on Advanced Video and Signal Based Surveillance (AVSS). IEEE, 2015, pp. 1–6.

[38] V. John, K. Yoneda, B. Qi, Z. Liu, and S. Mita, “Traffic light recognition in varying illumination using deep learning and saliency map,” in Proceedings of the International IEEE Conference on Intelligent Transportation Systems (ITSC), 2014, pp. 2286–2291.

[39] N.M. O’Boyle, “Towards a universal SMILES representation-a standard method to generate canonical SMILES based on the InChI,” Journal of Cheminformatics, vol. 4, no. 1, p. 22, 2012.

[40] S. Kim, P. A. Thiessen, E. E. Bolton, J. Chen, G. Fu, A. Gindulyte, L. Han, J. He, S. He, B. A. Shoemaker et al., “PubChem substance and compound databases,” Nucleic Acids Research, vol. 44, no. D1, pp. D1202–D1213, 2016.

[41] G. Landrum. RDKit: Open-source cheminformatics. [Online]. Available: http://www.rdkit.org

[42] B. Ramsundar, P. Eastman, P. Walters, V. Pande, K. Leswing, and Z. Wu, Deep Learning for the Life Sciences. O’Reilly Media, 2019.

[43] T. N. Kipf and M. Welling, “Semi-supervised classification with graph convolutional networks,” Proceedings of the International Conference on Learning Representations (ICLR), 2017.

[44] P. Velicković, G. Cucurull, A. Casanova, A. Romero, P. Lio, and Y. Bengio, “Graph attention networks,” Proceedings of the International Conference on Learning Representations (ICLR), 2018.

[45] K. Xu, W. Hu, J. Leskovec, and S. Jegelka, “How Powerful are Graph Neural Networks?” Proceedings of the International Conference on Learning Representations (ICLR), 2019.

[46] A. Krizhevsky, I. Sutskever, and G. E. Hinton, “Imagenet classification with deep convolutional neural networks,” in Advances in Neural Information Processing Systems, 2012, pp. 1097–1105.

[47] K. He, X. Zhang, S. Ren, and J. Sun, “Deep residual learning for image recognition,” in Proceedings of the IEEE Conference on Computer Vision and Pattern Recognition, 2016, pp. 770–778.

[48] K. Hornik, M. Stinchcombe, H. White et al., “Multilayer feedforward networks are universal approximators.” Neural Networks, vol. 2, no. 5, pp. 359–366, 1989.

[49] I. Sutskever, O. Vinyals, and Q. V. Le, “Sequence to sequence learning with neural networks,” in Advances in Neural Information Processing Systems, 2014, pp. 3104–3112.

[50] A. Vaswani, N. Shazeer, N. Parmar, J. Uszkoreit, L. Jones, A. N. Gomez, Ł. Kaiser, and I. Polosukhin, “Attention is all you need,” in Advances in Neural Information Processing Systems, 2017, pp. 5998– 6008.

[51] M. P. Menden, F. Iorio, M. Garnett, U. McDermott, C. H. Benes, P. J. Ballester, and J. Saez-Rodriguez, “Machine learning prediction of cancer cell sensitivity to drugs based on genomic and chemical properties,” PLoS ONE, vol. 8, no. 4, 2013.

[52] J. Codony-Servat, M. A. Tapia, M. Bosch, C. Oliva, J. Domingo-Domenech, B. Mellado, M. Rolfe, J. S. Ross, P. Gascon, A. Rovira et al., “Differential cellular and molecular effects of bortezomib, a proteasome inhibitor, in human breast cancer cells,” Molecular Cancer Therapeutics, vol. 5, no. 3, pp. 665–675, 2006.

[53] Friedman, Adam and Amzallag, Arnaud and Pruteanu-Malinici, Iulian and Baniya, Subash and Cooper, Zachary and Piris, Adriano and Hargreaves, Leeza and Igras, Vivien and Frederick, Dennie and Lawrence, Donald and Haber, Daniel and Flaherty, Keith and Wargo, Jennifer and Ramaswamy, Sridhar and Benes, Cyril and Fisher, David, “Landscape of targeted anti-cancer drug synergies in melanoma identifies a novel BRAF-VEGFR/PDGFR combination treatment,” PLOS ONE, vol. 10, p. e0140310. 10 2015.

[54] S. Rosenberg, V. T. DeVita, and S. Hellman, Cancer: Principles & Practice of Oncology. Lippincott Williams & Wilkins, 2005.

[55] J. M. Corton, J. G. Gillespie, S. A. Hawley, and D. G. Hardie, “5-Aminoimidazole-4-carboxamide ribonucleoside: a specific method for activating AMP-activated protein kinase in intact cells?” European Journal of Biochemistry, vol. 229, no. 2, pp. 558–565, 1995.

[56] B. Ricciuti, G. C. Leonardi, and M. Brambilla, “Emerging Biomarkers in the Era of Personalized Cancer Medicine,” Disease Markers, vol. 2019, 2019.

[57] A. C. Winters and K. M. Bernt, “MLL-rearranged leukemias—an update on science and clinical approaches,” Frontiers in Pediatrics, vol. 5, p. 4, 2017.

[58] N. Aben, D. Vis, M. Michaut, and L. Wessels, “Tandem: a two-stage approach to maximize interpretability of drug response models based on multiple molecular data types,” Bioinformatics, vol. 32, pp. i413–i420, 09 2016.

[59] J. Costello, L. Heiser, E. Georgii, M. Gönen, M. Menden, N. Wang, M. Bansal, M. Ammad-ud din, P. Hintsanen, S. Khan, J. Mpindi, O. Kallioniemi, A. Honkela, T. Aittokallio, K. Wennerberg, J. Collins, D. Gallahan, D. Singer, J. Saez-Rodriguez, and G. Van Westen, “A community effort to assess and improve drug sensitivity prediction algorithms,” Nature Biotechnology, 06 2014.

[60] I. S. Jang, E. Chaibub Neto, J. Guinney, S. Friend, and A. Margolin, “Systematic assessment of analytical methods for drug sensitivity prediction from cancer cell line data,” Pacific Symposium on Biocomputing. Pacific Symposium on Biocomputing, vol. 19, pp. 63–74, 01 2014.

